# Interhelical distance measurements support a hairpin conformation for Caveolin-1 in phospholipid bicelles

**DOI:** 10.1101/2025.07.08.663041

**Authors:** Katrina Brandmier, Soohyung Park, Wonpil Im, Kerney Jebrell Glover

## Abstract

Previous molecular dynamics (MD) simulations of caveolin-1 (Cav1) in our labs revealed the possibility of two stable conformations of its intramembrane helices (H1 and H2). To distinguish between these two conformations, experimental intramolecular distances obtained using FRET (Förster resonance energy transfer) were integrated into the MD simulations to better define the position of these helices. Two mutants of Cav1 were generated where an acceptor fluorophore, dansyl, was positioned at the N-terminal end and center of H1 (A87C, F99C respectively), while a donor fluorophore, a native tryptophan, was positioned at the C-terminal end of H2 (W128). For the A87C W128 mutant, a distance of 22.5 ± 0.3 Å was observed while a distance of 24.4 ± 0.2 Å was observed for the F99C W128 mutant in phospholipid bicelles. These experimental FRET distances were compared to distances in MD simulations of over 100 Cav1 structures. Together these studies support that H1 and H2 adopt a hairpin conformation in bicelles.

## Introduction

Caveolae are omega-shaped invaginations present in the plasma membrane of various cell types including adipocytes, endothelial cells, and fibroblasts. (1–3) Caveolae are responsible for key cellular functions including cell signaling, mechanoprotection, and endocytosis. (4–7) Found in high abundance in caveolae is the integral membrane protein caveolin, which has three highly-homologous tissue-specific isoforms (−1, -2, -3). (8–10) Both caveolin-1 (Cav1) and caveolin-3 (Cav3) have been shown to be necessary for the formation of caveolae, as the absence of the Cav1 or Cav3 gene results in the loss of caveolae from the plasma membrane. (11–13) Misregulation of caveolins has been linked to a wide variety of human disease states including heart and pulmonary disease, cancer, and muscular dystrophy. (14,15)

Caveolins are found on the intracellular leaflet of the plasma membrane as both the N- and C-termini are on the cytosolic side and there is no extracellular portion of the protein (16). This disposition along with the tight association of caveolin with the plasma membrane (i.e., not peripheral) has led to the postulation that caveolins possess an unusual intramembrane turn, although this has yet to be definitively established. (17,18) Cav1 is the most ubiquitous of the three isoforms and hence has been most thoroughly characterized structurally. Although a complete secondary structural characterization of full-length Cav1 is not yet available, a residue-by-residue secondary structure characterization for residues 62-178, a segment that comprises approximately 66% of the sequence, has been accomplished. (19) This study revealed that Cav1 contains three α-helices, dubbed Helix 1 (H1), Helix 2 (H2), and Helix 3 (H3) *(Figure 1A*). (19)) H1 and H2 are proposed to be the helices that form the intramembrane loop while H3 forms a long amphipathic helix that rests on the surface of the membrane. This is in agreement with the secondary structure analysis of Cav3 which showed a similar helical arrangement. (20) Cav1 also appears to have the ability to present itself in a variety of oligomeric forms; although, the oligomers observed seem to be highly dependent on parameters such as the detergent and/or method used to isolate the protein. (21–28) Furthermore, which, if any, of these oligomeric forms are representative of the actual state of Cav1 in caveolae in vivo remains enigmatic.

**Figure 1.**
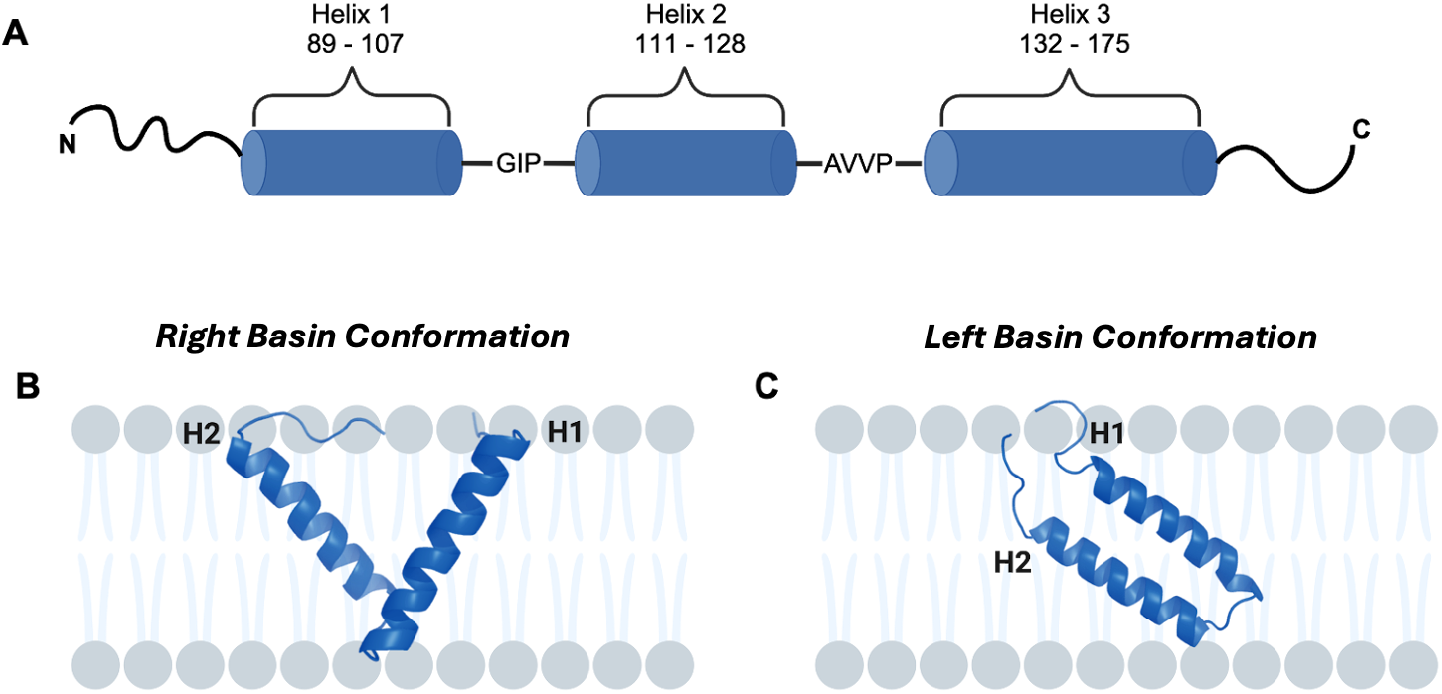
Cartoon representation of Cav1_62-178_ secondary structure **(A)**, Cav1_82-136_ right basin model **(B)**, and Cav1_82-136_ left basin model **(C)**.

Recently, we performed molecular dynamics (MD) and free energy simulations of H1 and H2 of Cav1 (residues 82-136) that revealed two roughly energetically equivalent conformations in a 1,2-dimyristol-*sn*-glycero-3-phosphocholine (DMPC) bilayer: the right basin (RB) and left basin (LB) (*Figure 1B, 1C)*. (29) Interestingly, the basis of stabilization is different. In the RB model H1 and H2 are stabilized by hydrogen bonds to the lipid headgroup regions whereas, in the LB model, H1 and H2 are stabilized by interhelical hydrogen bonds. (29) Additionally, the LB model shows a ∼19° angle between H1 and H2, while in the RB model, the angle between H1 and H2 is ∼76°. (29)

In order to further refine the molecular dynamics simulations, we employed steady-state Förster resonance energy transfer (FRET) measurements to determine key intramolecular distances. The FRET pair used was dansyl as the acceptor and tryptophan as the donor, a pair that has been utilized in FRET studies of other membrane proteins. (30) Advantageously, tryptophan is native to the protein, removing the challenges of labeling with two fluorophores in a site-specific way. These studies showed that the LB conformation is the most likely conformation of Cav1 (*Figure 1C*).

## Materials and Methods

Empigen BB^®^ was purchased from Sigma-Aldrich (St. Louis, MO). Ni Sepharose 6 Fast Flow resin and Sephacryl^®^ S300 HR 16/60 were purchased from Cytiva Life Sciences (Marlborough, MA). 1,5-IAEDANS was purchased from Setareh Biotech (Eugene, OR). 1-Naphthalenesulfonamide, 5-(dimethylamino)-N-[2-(2-pyridinyldithio)ethyl] (dansyl PDS) was synthesized in-house. Lysozyme was purchased from ThermoFisher Scientific (Waltham, MA). Tris(2-carboxyethyl)phosphine hydrochloride (TCEP) was purchased from GoldBio (St. Louis, MO). 96 well plates were purchased from Greiner Bio-One (Monroe, NC). Amicon^®^ Ultra centrifugal concentrators were purchased from MilliporeSigma (Burlington, MA). All other reagents were of standard ACS grade.

### Caveolin-1 FRET mutagenesis

Cav1 contains four native tryptophans (W85, W98, W115, and W128) and each tryptophan was explored as a single tryptophan mutant of Cav1. However, only W128 demonstrated monomeric behavior in Empigen BB^®^. Cav1 contains three cysteine residues which are natively palmitoylated, however studies have shown that palmitoylation is not required for proper trafficking of caveolae. Therefore these cysteine residues were mutated to serine (C133S, C143S, C156S) to avoid biologically irrelevant disulfide bonding. (31)

To design the FRET constructs, the following Cav1 constructs were made:

Construct 1 (A87C W128): W85F A87C W98F W115F C133S C143S C156S

Construct 2 (F99C W128): W85F W98F F99C W115F C133S C143S C156S

Construct 3 (W128): W85F W98F W115F C133S C143S C156S

All mutations were made using the QuikChange^®^ Site-Directed Mutagenesis Kit purchased from Agilent (Santa Clara, CA).

### Caveolin-1 expression and membrane preparation

Cav1 with a C-terminal myc tag and relevant mutations was cloned in the pET-24a(+) vector appending a C-terminal hexahistidine tag, and transformed into BL21(DE3) *E. coli* cells for overexpression using the autoinduction method by Studier *et a*l. (32) In short, 1000 µL of an overnight (20 h) starter culture in MDG media was used to inoculate 1 L of ZYM-5052 media in a 6 L Erlenmeyer flask. The culture was shaken at 25°C for 24 hours at 250 rpm. Cells were harvested by centrifugation at 8200 x g for 30 min at 4°C in 4 × 250 mL centrifugation bottles. Cells were then washed with 0.9% (w/v) NaCl recentrifuged, and stored at -20°C until ready for use. One 250 mL pellet was resuspended in 25 mL of 1x TAE (40 mM Tris-acetate, 1 mM EDTA, pH 8.0) supplemented with 1 mg/mL lysozyme and stirred at 4°C for 1 hour, then sonicated using a Branson sonifier 450 equipped with a flat tip (duty cycle 3, output power 3) until the cells were fully homogenized (∼15 min). The temperature was kept below 10°C during the sonication process. The homogenized cells were then centrifuged at 7000 x g for 30 minutes to pellet cellular debris. Carefully, the supernatant was removed and the membranes pelleted by centrifugation at 230,000 x g for 20 minutes at 4°C. Membranes were then washed by re-homogenizing each membrane in 6 mL of 1 × TAE using a 3 mL syringe and 21 gauge x 1 (0.8mm x 25mm) needle. 0.5 mM TCEP was present for all cysteine-containing constructs. Washed membranes were re-pelleted by centrifugation at 230,000 x *g* for 20 minutes at 4°C.

### Caveolin-1 purification and reconstitution into bicelles

Washed membranes were resuspended in 4 mL of 50 mM sodium phosphate pH 8.0, 500 mM sodium chloride, 2% (v/v) Empigen BB^®^, and 0.5 mM TCEP + 1mM dansyl PDS for the constructs and 1 mM 1,5 - IAEDANS 2 (construct 3 does not require labeling). Cav1 samples were purified and reconstituted into bicelles as previously described. (33)

### Determination of Forster Distance for Trp-Dansyl Pair

The Förster distance, *R*_0_ depends on the quantum yield of the donor (*Q*_*D*_), the degree of spectral overlap between the donor emission and acceptor absorption (*J*), the refractive index of the solution (*n*), and the relative orientation in space (*k*^2^), following the equation:

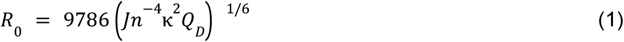

The quantum yield of W128 was determined using a relative method (34). Quinine sulfate in 0.1 M H_2_SO_4_ was selected as the reference at 25°C. The quantum yield of W128 was determined to be 0.16 ± 0.01. The degree of spectral overlap was calculated as 5.839 × 10^13^ M^-1^ cm^-1^nm^4^. The relative refractive index of the solution was measured as 1.3362 using an Anton Paar Abbemat 3200 benchtop refractometer. Assumed values of ⅔ for *k*^2^ were used. The *R*_0_ for the W128 - dansyl pair was calculated as 22.1Å.

### Fraction Labeled of Donor and Acceptor Constructs

Fraction labeled was determined using Equation 2, where A_337_ and A_280_ represent the absorbance of the sample corrected at 337 nm and 280 nm, respectively. *ε*_*D*337_ represents the extinction coefficient of dansyl at 337 nm which is 5,700 cm^-1^M^-1^. ε_*p*280_ is the extinction coefficient of the protein, 18,910 cm^-1^M^-1^.

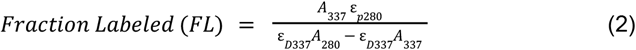

### Fluorescence Intensity Measurements

Concentrations of protein pairs were matched using a micro BCA assay. 150 µL of equal concentrations of the donor sample and acceptor sample were placed in a 96 well plate alongside reference samples. Samples were excited at 295 nm and emission was collected at 345 nm, with 100 flashes using an Infinite M Nano Tecan plate reader. Corrected fluorescence intensities were utilized for FRET calculations. All measurements were performed in triplicate.

### Determination of Intramolecular Distances

FRET efficiency (E) can be calculated using the equation below:

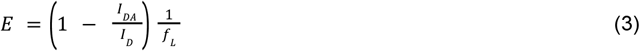

where I_DA_ and I_D_ are the fluorescence intensity in the presence and absence of the acceptor, respectively. f_L_ is the fraction labeled calculated in Equation 1.

Following FRET efficiency, intramolecular distances can be calculated using the equation below:

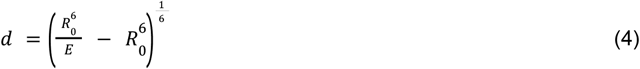

where R_0_ is the Förster distance and E is the FRET efficiency. The Förster distance for this fluorophore pair is 22.1 Å. (30)

### Molecular Dynamic Simulations

To simulate FRET experiments, we sampled the distances between fluorophore pairs for the RB and LB Cav1 constructs, whose conformations were chosen from our previous simulations in explicit membranes. (35) Note that helix 3 was included in these simulations and also in the donor-acceptor distance calculations. The constructs simulated were exact mimics of the constructs prepared in the experimental FRET Studies (see section titled *Caveolin-1 FRET mutagenesis*).Using a method analogous to DEER Spin-Pair Distributor (36) in CHARMM-GUI (37), donor-acceptor distances were sampled from simulations of Cav1 in vacuum. In these simulations, the Cav1 structure was fixed, except the fluorophore pair, allowing us to obtain a distance distribution between the pair for a given Cav1 conformation. For each Cav1 structure, an initial acceptor conformation was chosen by exploring χ dihedral angles along the sidechain of the mutated Cys residue. For the first three χ dihedrals, eight combinations were considered, based on those for the methanethiosulfonate spin label. (38) For the outer χ dihedrals, gauche (± 60°) and trans (180°) dihedrals were considered, except the last χ. Due to the bulky aromatic ring, the last χ was searched among ±90° and 0°. The lowest energy conformation was chosen for the subsequent simulations.

A 10-ns Langevin dynamics simulation was performed using CHARMM with the C36 force field (39) and CGenFF for the acceptor. (40) The van der Waals and electrostatic interactions were calculated using those in the DEER Spin-Pair Distributor input script, with the friction coefficient of 5.0 ps^-1^. The simulation was run at a temperature of 1,000 K for better sampling of fluorophore conformations. An integration time step of 1 fs was used, and trajectory was saved every 1 ps. To account for Cav1 dynamics, we selected 100 structures for both the LB and RB conformations for each Cav1 construct from our previous simulations. (35) From the trajectories, distances between the centers of mass of aromatic rings *(Figure 2A, 2B)* were calculated. Although we used only dansyl fluorophore in simulations, distance between the fluorophore pair in A87C W128 would not be significantly affected due to its comparable size to 1,5-IAEDANS (*Figure 2A, 2B*).

**Figure 2.**
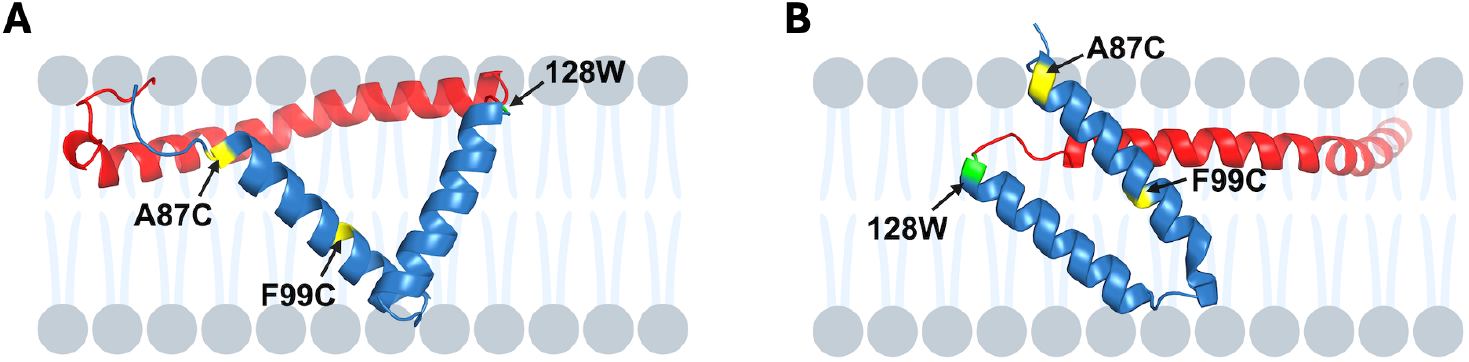
Centroid structures of Cav1 RB ***(A)*** and LB ***(B)***. *Yellow* represents the acceptor positions, A87C and F99C, and *green* represents the donor position, W128. H1 and H2 are shown in *blue* while H3 is shown in *red*.

## Results and Discussion

To better elucidate the tertiary structure of Cav1, experimental FRET-derived distances were used to examine which conformation, LB or RB, is most likely. FRET is a phenomenon where energy can be transferred between a donor fluorophore and an acceptor fluorophore, as long as the excitation of the acceptor fluorophore overlaps spectrally with the emission of the donor fluorophore. FRET is highly distance-dependent and can measure distances between 10 Å and 100 Å (41). The FRET pair that we chose was dansyl and tryptophan. Adventitiously, the tryptophan is native to Cav1 so the challenge of site-specifically labeling the protein with both donor and acceptor was averted.

FRET pairs were initially selected based on predicted C*α* - C*α* distances that differed between the LB and RB conformation. Table 1 shows predicted C*α* - C*α* distances for four possible FRET pairs, that result in distinguishable distances between the RB and LB conformations. Following mutagenesis and purification of each pair, those with W98 as the donor (I121C 98W and L125C 98W) exhibited high levels of dimer, which would result in *inter*molecular distances. As a result, we proceeded further experimental analysis with A87C W128 and F99C W128 which both exhibited monomeric behavior.

**Table 1.** Comparison of predicted C*α* - C*α* distances between FRET donor - acceptor pairs for three Cav-1 structural models and experimental de novo FRET distance in DMPC-DHPC bicelles. The right and left basin models are representative conformations from our previous work (29). A chain from the cryo-EM structure model is denoted as its PDB ID, 72C0 (24).

| Donor-Acceptor<br>FRET pair | $C\alpha$ - $C\alpha$ Distance (Å) | | | Experimental<br>FRET Distance<br>(Å) |
| --- | --- | --- | --- | --- |
|  | Right Basin<br>Conformation | Left Basin<br>Conformation | PDB ID: 7SC0 |  |
| A87C W128 | 35.5 | 10.7 | 61.1 | 22.5 ± 0.3 |
| F99C W128 | 26.0 | 15.5 | 45.0 | 22.4 ± 0.2 |
| I121C 98W | 19.8 | 8.8 | - | - |
| L125C 98W | 24.3 | 11.8 | - | - |

**Table 2.** FRET efficiency and calculated donor - acceptor distance for the A87C W128 mutant. Reported values represent the mean ± SEM.

| Caveolin-1 Mutant | FRET Efficiency | Calculated Distance (Å) |
| --- | --- | --- |
| A87C W128 | $0.387 \pm 0.017$ | $22.5 \pm 0.3$ |

**Table 3.** FRET efficiency and calculated donor - acceptor distance for the F99C W128 mutant. Reported values represent the mean ± SEM. M.

| Caveolin-1 Mutant | FRET Efficiency | Calculated Distance (Å) |
| --- | --- | --- |
| F99C W128 | $0.347 \pm 0.008$ | $24.4 \pm 0.2$ |

The first construct was A87C W128, where A87 was mutated to cysteine for labeling with the dansyl acceptor fluorophore. This pair probes the distance between the head of H1 (position 87) and the tail of H2 (position 128). The second construct, F99C 128W, F99 was mutated to cysteine for labeling with the dansyl acceptor fluorophore and resides in the center of H1, while W128 was kept as the donor site. Figure 2 shows the FRET pairs on the LB and RB conformations.

### Intramolecular distance between A87C W128 FRET in phospholipid bicelles

Experimentally, the distance between A87C and W128 was determined to be 22.5 ± 0.3 Å by FRET in phospholipid bicelles (Table 1). Given that the acceptor fluorophore is positioned at the N-terminal end of H1 (A87C), and the donor is positioned at the C-terminal end of H2, 42 amino acids separate them. Based on previous NMR studies from our lab, of those 42 residues only three, **GIP**, are excluded from **α**-helical structure (19,42). Therefore, if H1 and H2 are ideal **α**-helices extended linearly, the distance between the donor and the acceptor fluorophore would be approximately 58.5 Å. Clearly, a distance of 58.5 Å, is much greater than 2 times R_0_ (41.8 Å) and would produce negligible levels of FRET. Therefore, we can rule out an extended linear conformation of H1 and H2.

Next, simulations were run of both the LB and RB conformations with the dansyl (acceptor) fluorophore attached at position 87. Figure 3A shows the distance distributions between the donor and acceptor for the 100 structures of LB and RB conformations, which are well separated without any overlap. The vertical line represents the experimentally determined FRET distance of 22.5 ± 0.3 Å, and it is clear that it overlaps with the LB structures (red) much better than the RB structures (blue). Figure 3B shows a representative structure for LB conformation, highlighting the donor and acceptor positions. If Cav1 preferred the RB conformation, very little FRET would be observed as most of the RB conformations are greater than 2 times R_0_ (41.8 Å) which would result in FRET efficiencies of less than 1.5%. This clearly supports that Cav1 agrees much better with the LB conformation, placing H1 and H2 at a smaller angle with respect to each other.

**Figure 3.**
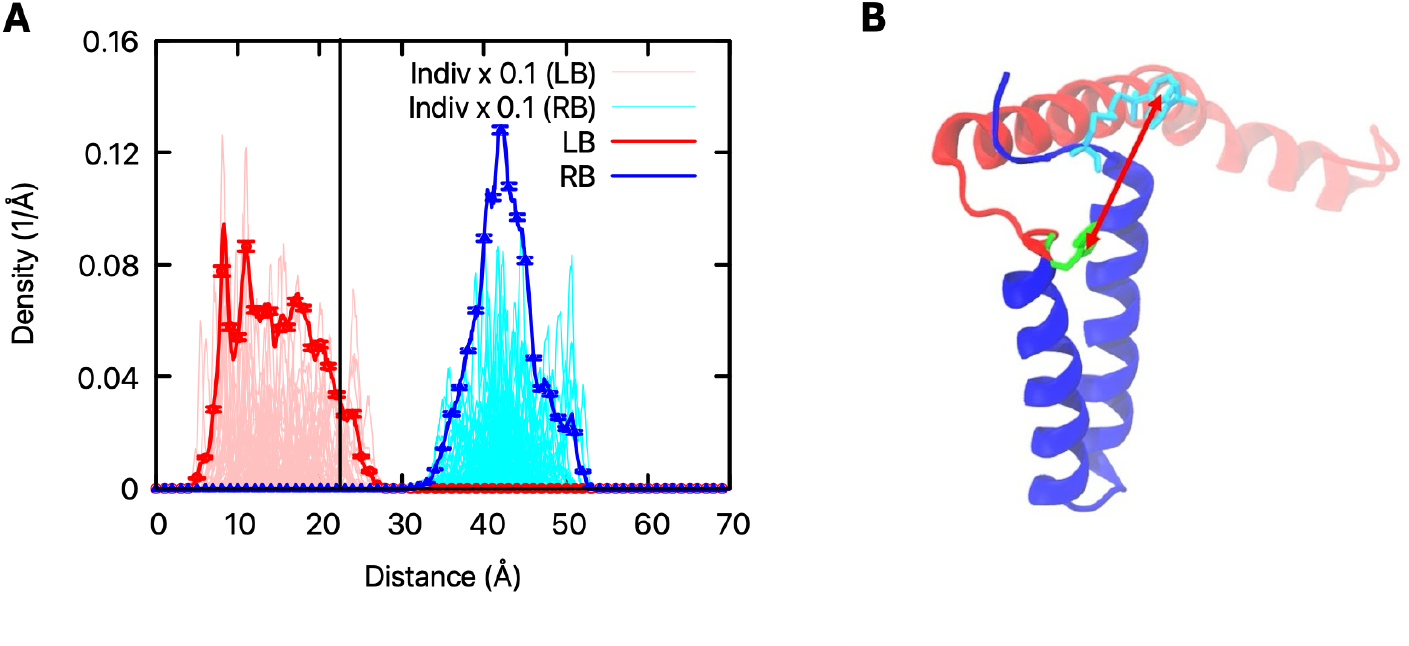
***(A)*** Average distance distribution between dansyl at position 87 and tryptophan at position 128 across 100 structures of Cav1 for both the LB (red) and RB (blue) conformations. Shown in *pink* (LB) and *cyan* (RB) are the distances for each 100 Cav1 structures. The vertical black line represents the experimentally determined FRET distance of 22.5 ± 0.3 Å in bicelles. ***(B)*** A representative structure of the LB conformations with dansyl highlighted in *cyan* and tryptophan highlighted in *green* H1 and H2 are *blue* and H3 is *red*.

### Intramolecular distance between F99C W128 FRET in bicelles

The experimental distance between F99C W128 calculated through FRET was determined to be 24.4 ± 0.2 Å in DMPC-DHPC bicelles. The donor-acceptor distance distributions for both the LB and RB conformations from simulations showed substantial overlap (Figure 4A). Differently from expected C*α* - C*α* distances (Table 1), the experimental distance remains compatible with both conformations when fluorophore rotamers are considered and limits the discriminative power of the F99C W128 pair alone. These findings highlight the importance of careful consideration of fluorophore rotamers in the interpretation of FRET distances, which can be significantly different from simple C*α* - C*α* distances. Additionally, Cav1 presents unique difficulties due to the high degree of sequence conservation. Many residues in the transmembrane domain appear to be functionally or structurally critical. Thus, limiting the range of viable mutations, making it challenging to identify FRET pairs that are able to distinguish between the RB and LB conformation and are biochemically well-behaved. Nevertheless, the pairs that do satisfy these constraints consistently reinforce a compact, rather than extended, arrangement supporting the physiological relevance of this conformation in phospholipid bicelles.

**Figure 4.**
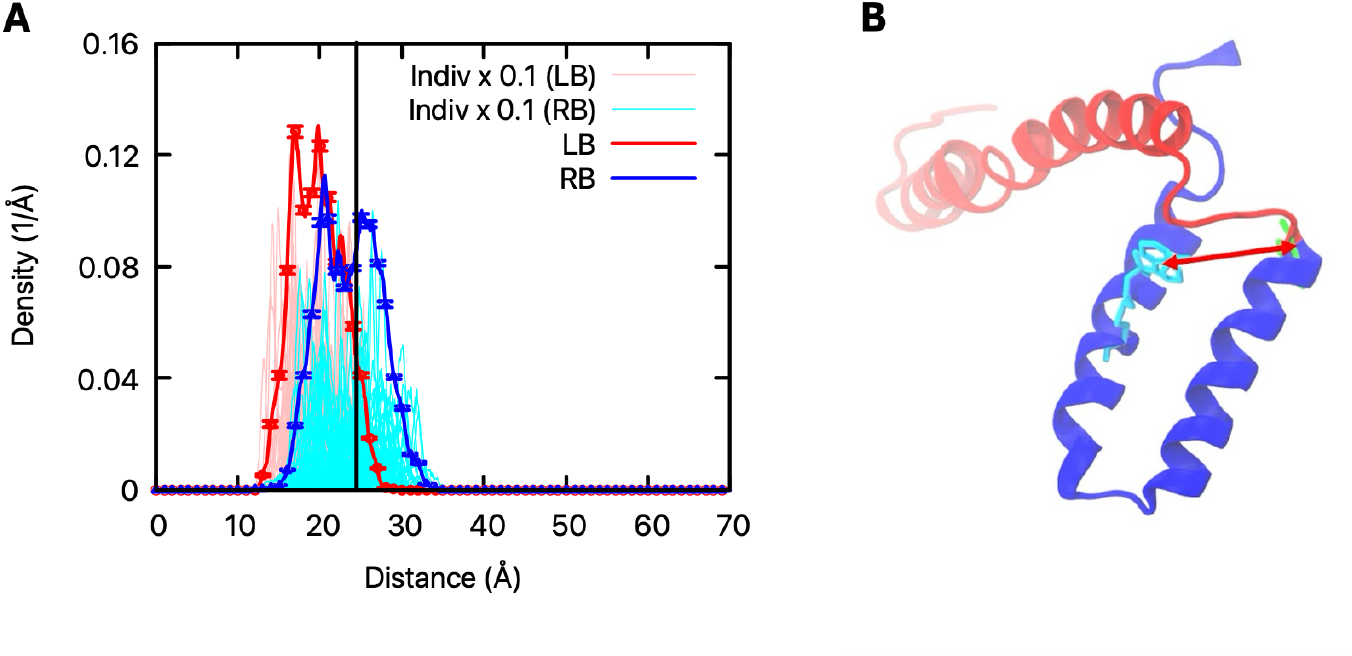
***(A)*** Average distance distribution between dansyl at position 99 and tryptophan at position 128 across 100 structures of Cav1 for both the LB (red) and RB (blue) conformations. Shown in *pink* (LB) and *cyan* (RB) are the distances for each 100 Cav1 structures. The vertical black line represents the experimental distance of 24.4 Å ± 0.2 Å. ***(B)*** A representative structure of the LB confirmation with dansyl highlighted in *cyan* and tryptophan highlighted *green* H1 and H2 are *blue* and H3 is *red*.

Recently, a cryo-electron microscopy (cryo-EM) structure of Cav1 revealed the organization of a detergent-stabilized oligomeric complex, termed the 8S complex (PDB 72C0). This structure was determined from recombinant human Cav1 expressed in E. coli and purified in n-dodcel-B-D-maltoside (DDM) detergent, capturing an oligomer composed of 11 protomers arranged in a disc-like assembly. Each monomer adopts a highly extended conformation, with long intramembrane helices oriented nearly parallel to the membrane plane (24). To contextualize our de novo FRET distances, we calculated the C*α* - C*α* distance for each of our FRET pairs using a single monomeric subunit extracted from the 8S Cav1 structure. The predicted distances for the 72C0 model were substantially longer than both those measured experimentally and in the RB and LB C*α* - C*α* measurements (Table 1). Importantly, the predicted distances were greater than 2R_0_ (44.2Å) away, placing them outside of the range for measurable FRET efficiency. Given that we observed significant FRET efficiency with both pairs, the conformation of Cav1 experimentally observed here must adopt a hairpin conformation unlike the monomeric subunit of the 8S model.

Taken together, our experimental FRET distances, supported by MD simulations, are consistent with the left basin conformation of Cav1 and strongly favor Cav1 adopting an intramembrane turn in DMPC-DHPC bicelles. Furthermore, comparison to the monomeric subunit of the 8S Cav1 cryo-EM structure reveals predicted distances that are incompatible with the observed FRET efficiencies, ruling out a fully extended model. Collectively, these findings support a compact hairpin conformation for Cav1 consistent with the left basin model and provides further insight into the structural arrangement of Cav1 in a membrane mimic.

## Acknowledgements

Figure 1 and 2 were created using BioRender.com. This work was supported by the NIH R15 GM141606-01 awarded to Kerney Jebrell Glover and Wonpil Im. The authors thank Caroline McDowell and Rosa Medina for computational and laboratory assistance, respectively. Additionally, the authors thank Christophe Guillon and Jaya Saxena for the use of their Infinite M Nano Tecan plate reader.

